# Nanotraps for the containment and clearance of SARS-CoV-2

**DOI:** 10.1101/2021.02.01.428871

**Authors:** Min Chen, Jillian Rosenberg, Xiaolei Cai, Andy Chao Hsuan Lee, Jiuyun Shi, Mindy Nguyen, Thirushan Wignakumar, Vikranth Mirle, Arianna Joy Edobor, John Fung, Jessica Scott Donington, Kumaran Shanmugarajah, Eugene Chang, Glenn Randall, Pablo Penaloza-MacMaster, Bozhi Tian, Maria Lucia Madariaga, Jun Huang

## Abstract

SARS-CoV-2 enters host cells through its viral spike protein binding to angiotensin-converting enzyme 2 (ACE2) receptors on the host cells. Here we show functionalized nanoparticles, termed “Nanotraps”, completely inhibited SARS-CoV-2 infection by blocking the interaction between the spike protein of SARS-CoV-2 and the ACE2 of host cells. The liposomal-based Nanotrap surfaces were functionalized with either recombinant ACE2 proteins or anti-SARS-CoV-2 neutralizing antibodies and phagocytosis-specific phosphatidylserines. The Nanotraps effectively captured SARS-CoV-2 and completely blocked SARS-CoV-2 infection to ACE2-expressing human cell lines and primary lung cells; the phosphatidylserine triggered subsequent phagocytosis of the virus-bound, biodegradable Nanotraps by macrophages, leading to the clearance of pseudotyped and authentic virus *in vitro*. Furthermore, the Nanotraps demonstrated excellent biosafety profile *in vitro* and *in vivo*. Finally, the Nanotraps inhibited pseudotyped SARS-CoV-2 infection in live human lungs in an *ex vivo* lung perfusion system. In summary, Nanotraps represent a new nanomedicine for the inhibition of SARS-CoV-2 infection.

**Highlights:**

- Nanotraps block interaction between SARS-CoV-2 spike protein and host ACE2 receptors
- Nanotraps trigger macrophages to engulf and clear virus without becoming infected
- Nanotraps showed excellent biosafety profiles *in vitro* and *in vivo*
- Nanotraps blocked infection to living human lungs in *ex vivo* lung perfusion system

**Progress and Potential:** To address the global challenge of creating treatments for SARS-CoV-2 infection, we devised a nanomedicine termed “Nanotraps” that can completely capture and eliminate the SARS-CoV-2 virus. The Nanotraps integrate protein engineering, immunology, and nanotechnology and are effective, biocompatible, safe, stable, feasible for mass production. The Nanotraps have the potential to be formulated into a nasal spray or inhaler for easy administration and direct delivery to the respiratory system, or as an oral or ocular liquid, or subcutaneous, intramuscular or intravenous injection to target different sites of SARS-CoV-2 exposure, thus offering flexibility in administration and treatment. More broadly, the highly versatile Nanotrap platform could be further developed into new vaccines and therapeutics against a broad range of diseases in infection, autoimmunity and cancer, by incorporating with different small molecule drugs, RNA, DNA, peptides, recombinant proteins, and antibodies.

## Introduction

The severe acute respiratory syndrome coronavirus 2 (SARS-CoV-2) caused the global pandemic of coronavirus disease 2019 (COVID-19). As of January 8, 2021, SARS-CoV-2 has spread to over 180 countries and has resulted in more than 88.2 million infections and over 1.9million deaths globally(Dong et al., 2020). Despite tremendous efforts devoted to drug development, safe and effective medicines to treat SARS-CoV-2 infection are largely lacking. Given that the virus is within nanoscale, nanomaterial based delivery systems are expected to play a paramount role in the success of prophylactic or therapeutic approaches(Florindo et al., 2020; Shin et al., 2020). To combat this highly contagious virus, here we set out to devise a nanomedicine termed “Nanotraps” to inhibit SARS-CoV-2 infection.

To gain entry to host cells for infection, SARS-CoV-2 surface spike protein binds to its receptor human angiotensin-converting enzyme 2 (ACE2) with high affinity(Shang et al., 2020b, 2020a; Vabret et al., 2020; Walls et al., 2020). Blocking entry of SARS-CoV-2 to host cells is one of the most effective ways to prevent infection. To achieve this goal, both soluble recombinant ACE2 proteins(Lei et al., 2020; Monteil et al., 2020) and anti-SARS-CoV-2 neutralizing antibodies(Cao et al., 2020; Chen et al., 2020b; Chi et al., 2020; Rogers et al., 2020; Shi et al., 2020) have been developed to inhibit the interaction between SARS-CoV-2 spike protein and cell-surface ACE2, although they show limited potency(Chen et al., 2020a; Lei et al., 2020; Monteil et al., 2020). Inspired by tumor cells secreting PD-L1 exosomes to attenuate T-cell effector functions(Daassi et al., 2020), here we designed a therapeutic nanoparticle termed “Nanotrap” to inhibit SARS-CoV-2 infection. The Nanotrap surfaces were functionalized with either ACE2 recombinant proteins or anti-SARS-CoV-2 neutralizing antibodies with high surface density. This design endowed the Nanotraps with high-avidity to outperform its soluble ACE2 or antibody counterparts to capture and contain SARS-CoV-2. Thus, the high avidity, small size, and high diffusivity of our newly engineered Nanotraps efficiently blocked the binding of SARS-CoV-2 to ACE2-expressing host cells including epithelial cells in the respiratory system, resulting in abrogation of SARS-CoV-2 entry.

Furthermore, we aimed to clear the virus after containment by the Nanotrap-mediated macrophage phagocytosis. The role of macrophages in the control of infections has long been documented(Gordon, 2016), and recent single-cell RNA sequencing found abundant monocyte-derived macrophages in the bronchoalveolar lavage fluid of COVID-19 patients(Liao et al., 2020). As professional phagocytes, macrophages engulf apoptotic cells by recognizing phosphatidylserine on the outer leaflet of the plasma membrane of apoptotic cells(Aderem and Underhill, 1999; Fadok et al., 1998; Wu et al., 2006). Phosphatidylserine coatings have been previously employed to enhance the uptake of liposomal nanoparticles by macrophages(Johnstone et al., 2001; Shah et al., 2019). Thus, our Nanotraps were further designed to guide the phosphatidylserine-specific phagocytosis by macrophages, enabling not only the containment but also the clearance of SARS-CoV-2 through binding and subsequent phagocytosis.

Herein, we engineered Nanotraps composed of a Food and Drug Administration (FDA)-approved polylactic acid (PLA) polymeric core, a liposome shell, surface ACE2/neutralizing antibodies, and phosphatidylserine ligands. Our Nanotraps completely blocked pseudotyped SARS-CoV-2 entry into susceptible ACE2-overexpressing HEK293T cells, lung epithelial A549 cells, and human primary lung cells, as well as authentic SARS-CoV-2 infection of Vero E6 cells. Subsequently, macrophages efficiently engulfed and neutralized virus-bound, biodegradable Nanotraps through phosphatidylserine-guided phagocytosis without causing infection to macrophages *in vitro*. Furthermore, *in vitro* cell culture and *in vivo* intratracheal administration of Nanotraps to immunocompetent mice demonstrated an excellent biosafety profile. Lastly, the Nanotraps completely inhibited infection of SARS-CoV-2 pseudovirus in live human lungs maintained under normothermic physiologic conditions on a clinically applicable *ex vivo* lung perfusion (EVLP) system(Cypel et al., 2008; Divithotawela et al., 2019), confirming the therapeutic efficacy. Our Nanotraps are safe, effective, biocompatible, ready for mass production, and convenient to use. It presents a new type of nanomedicine to effectively contain and clear SARS-CoV-2 for the prevention and treatment of COVID-19.

## RESULTS

### Design, synthesis, and characterization of Nanotraps

SARS-CoV-2 gains entry into host cells via surface spike proteins that bind to human ACE2 receptors on host cells with very high affinity(Hwang et al., 2020; Lan et al., 2020; Shang et al., 2020a; Wang et al., 2020a). To inhibit SARS-CoV-2 infection, we set out to engineer a family of nano-enabled virus-trapping particles, termed “Nanotraps”, to contain and clear SARS-CoV-2 (Figure 1A). We used an FDA-approved, biodegradable PLA polymeric core and liposome shell materials to synthesize the Nanotraps. Nanotraps with different diameters (200, 500, and 1200 nm) were synthesized by varying polymer concentrations (see Methods for details) (Figure S1A). The solid PLA core acts as a ‘cytoskeleton’ to provide mechanical stability, controlled morphology, biodegradability, and large surface area for nanoscale membrane coating and surface modification. The lipid shell enveloping the PLA core exhibits behavior similar to that of cell membranes. The lipid shell provides a nanoscopic platform and can interact with a wide variety of molecules(Allen and Cullis, 2013; Riley et al., 2019; Torchilin, 2005) either within the membrane or on the surface(Suk et al., 2016; Torchilin, 2014). Thus, we aimed to functionalize the Nanotrap surface with a molecular bait (a recombinant ACE2 protein or an anti-SARS-CoV-2 neutralizing antibody) and a phagocytosis-inducing ligand (phosphatidylserine). We hypothesized that (1) the high-density ACE2 or neutralizing antibodies on the Nanotraps can outcompete low-expression ACE2 on host cells in capturing SARS-CoV-2, thus enabling selective virus containment by the Nanotraps, and that (2) surface phosphatidylserine ligands on suitably sized Nanotraps can trigger subsequent phosphatidylserine-mediated phagocytosis by professional phagocytes, such as macrophages, thus enabling viral clearance (Figure 1A). The resultant structures were monodispersed and significantly smaller than mammalian cells, yet still large enough to bind several SARS-CoV-2 virions (Figure 1B-F).

**Figure 1:**
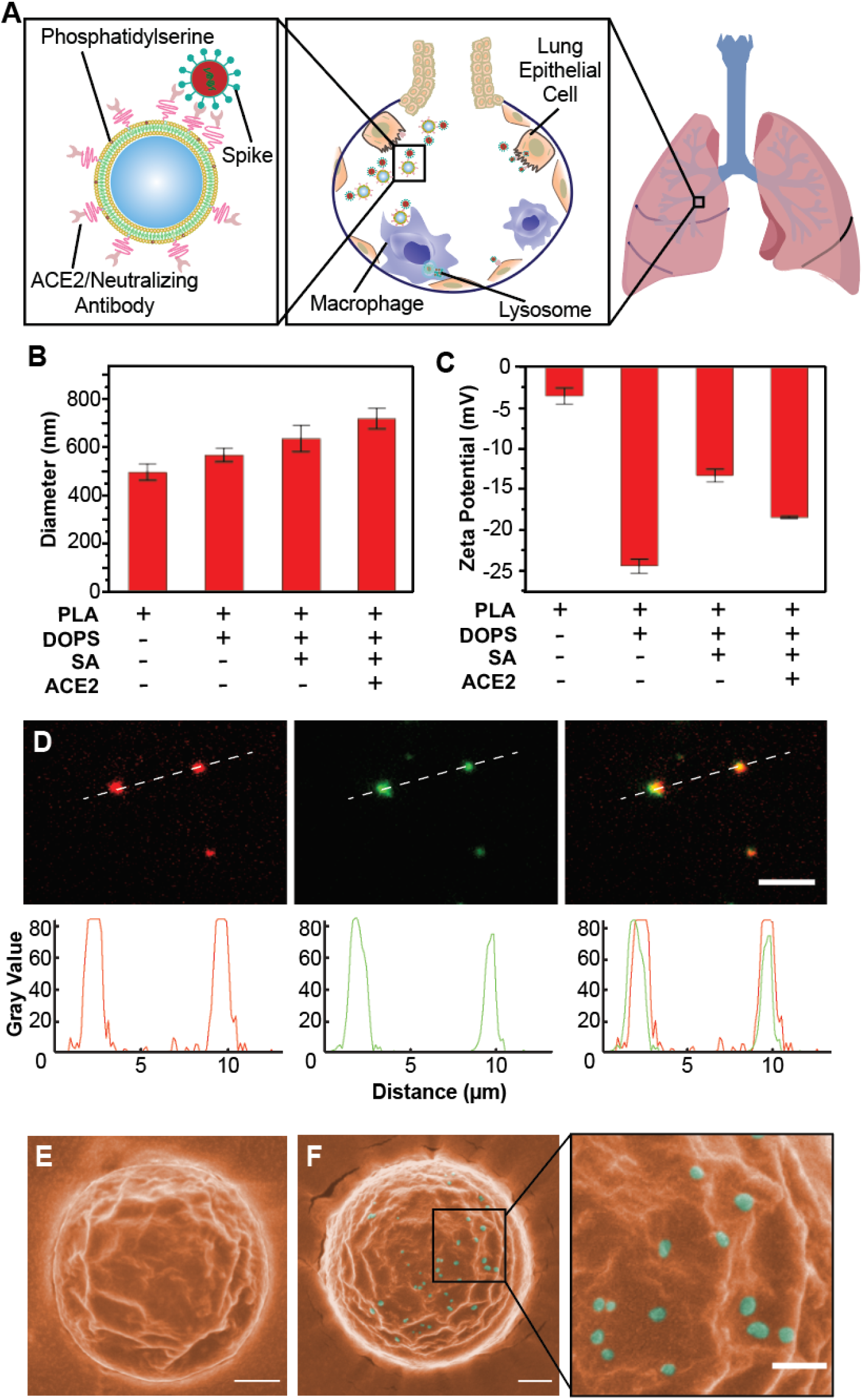
Schematic design, synthesis, and characterization of Nanotraps for SARS-CoV-2. (**A**) Schematic illustration showing the process of the Nanotraps with polymeric core coated with lipid-bilayer functionalized with ACE2 protein/neutralizing antibody. Following intratracheal administration, Nanotraps efficiently accumulated and trapped SARS-CoV-2 virionsin the lung tissue forming virus-Nanotrap complexes, which can be cleared by macrophages via phagocytosis, thereby blocking viral cell-entry. (**B-C**) Dynamic light scattering (B) and Zeta-potential measurements (C) during different stages of Nanotrap preparation. (**D**) Fluorescent images of the prepared Nanotraps with PLA polymeric core (DiD, red) and ACE2 (anti-ACE2-AF488, green). Scale bar represents 5 μm. Dotted lines represent displayed plot profile below. (**E-F**) Pseudocolored SEM images of Nanotraps alone (E, orange) or with SARS-CoV-2 pseudovirus (F, cyan). To better visualize the selectivity for viral binding, larger Nanotraps were imaged. Scale bar represents 300 nm.

To characterize the Nanotraps, we first used dynamic light scattering to measure the size dispersity of the constituent nanoparticles. Controlling particle size is important for tuning the phagocytosis efficacy, reproducible mechanical characteristics, and material biocompatibility(Champion et al., 2008; He et al., 2010). The hydrolyzed diameter of the Nanotraps was measured by dynamic light scattering, which increased with the addition of each molecule (Figure 1B). The zeta potential, which reflects the surface charges of the Nanotraps(Doane et al., 2012), was found to change slightly with the addition of each molecule to the Nanotrap surface (Figure 1C). We next used fluorescent labeling and total internal reflection fluorescence microscopy (TIRFM) to confirm the presence of the ACE2 moiety on the Nanotraps. The PLA polymeric core of each Nanotrap was labelled with a lipophilic carbocyanine dye: 1,1’-dioctadecyl-3,3,3’,3’- tetramethylindodicarbocyanine, 4-chlorobenzenesulfonate salt (DiD, red). The Nanotrap-ACE2 were further stained with an anti-ACE2 antibody labelled with Alexa fluor-488 dye (AF488, green). The TIRFM images clearly showed excellent co-localization between the Nanotrap core and surface ACE2 at the single particle level, confirmed by line scans of the fluorescent channels corresponding to each component (Figure 1D). These results demonstrated that we have successfully functionalized the Nanotraps with recombinant ACE2 protein. Finally, we employed scanning electron microscopy (SEM) to image the Nanotraps at the sub-nanometer level. SEM images showed that the Nanotraps were spherical and well-dispersed (Figure S1B). The Nanotraps appeared slightly crenellated in the SEM images, as the lipid layer may have shrunk due to the drying sample preparation procedure before imaging (Figure 1E). As expected, after incubation with the pseudotyped SARS-CoV-2 for 1 hour at 37 °C, the Nanotraps effectively captured the virus as evidenced by single virions clearly visualized on the surface of a Nanotrap; no freestanding virions were observed outside of the Nanotrap (Figure 1F).

### Phagocytosis of Nanotraps by macrophages

Macrophages are a class of phagocytes that engulf and clear cell debris, pathogens, microbes, cancer cells and other foreign intruders(Gordon, 2016). Macrophages specialize in the removal of dying or dead cells by recognizing phosphatidylserine on the outer leaflet of the plasma membrane of apoptotic cells(Aderem and Underhill, 1999; Fadok et al., 1998; Wu et al., 2006). Phosphatidylserine coatings have been previously employed to enhance macrophage uptake of liposomal nanoparticles(Han et al., 2016; Harel-Adar et al., 2011; Toita et al., 2016). Because phagocytosis by macrophages is highly dependent on size and surface phosphatidylserine, we determined the optimal size and surface phosphatidylserine density of Nanotraps. We first synthesized Nanotraps labelled with 3,3′-Dioctadecyloxacarbocyanine perchlorate (DiO) fluorescent dye on the PLA polymeric core with varying diameters: 200, 500, and 1200 nm (Figure S2A). After incubating Nanotraps with differentiated THP-1 (dTHP-1) macrophages(Lund et al., 2016) for varying time durations, we examined the size-dependent phagocytosis by dTHP-1 macrophages using flow cytometry. We found that the 500-nm Nanotraps outperformed the 200-nm and 1200-nm counterparts after 24- and 48-hour incubations, as demonstrated by the percent uptake (Figure 2A) and the mean fluorescent intensity of DiO dye (Figure 2A and Figure S2) in dTHP-1 macrophages. Accordingly, 500-nm PLA-core Nanotraps were chosen for all further experiments.

**Figure 2:**
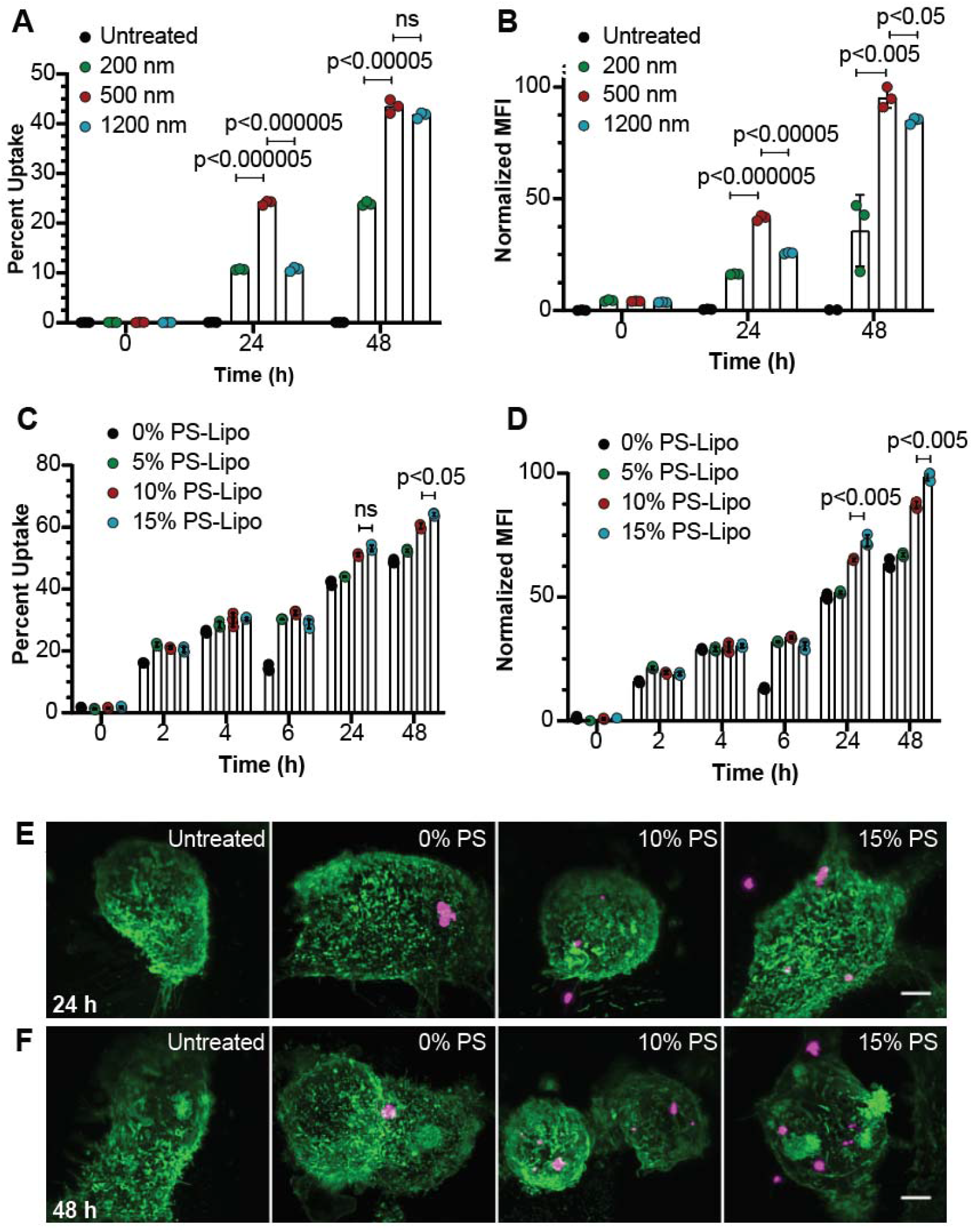
Phagocytosis of Nanotraps by macrophages. Nanotraps of varying size and phosphatidylserine (PS) densities were incubated with dTHP-1 macrophages. (**A-B**) Percent uptake (A) and mean fluorescent intensity (B) quantification of flow cytometry measurement of the internalization of Nanotraps with varying diameters. (**C-D**) Percent uptake (C) and mean fluorescent intensity (D) quantification of flow cytometry measurements of the internalization of Nanotraps with varying PS ratios. Data are shown as mean ± SD; unpaired t tests were conducted from three replicates. (**E-F**) Lattice light-sheet microscopy images of macrophages (Wheat Germ Agglutinin (WGA)-CF488, green) phagocytosing Nanotraps (DiD, magenta) after 24 hours (E) or 48 hours (F). Scale bar represents 5 μm.

We next functionalized the Nanotraps with varying surface densities of phosphatidylserine to induce phagocytosis by macrophages. The overall negative charge of the Nanotraps increases as the percentage of phosphatidylserine ratio increases, confirming the presence of phosphatidylserine (Figure S2B). We found that phagocytosis by macrophage was roughly correlated to the percentage of phosphatidylserine (Figure 2C-D). The phosphatidylserine-dependent phagocytosis of Nanotraps by dTHP-1 macrophages were further demonstrated by three-dimensional lattice light-sheet microscopy images (Figure 2E-F and Supplemental Video 1). These experiments not only confirmed successful functionalization of the Nanotraps, but also identified the optimal surface density of phosphatidylserine on the Nanotrap surfaces. Thus, we utilize the 500-nm core size and 15% surface phosphatidylserine Nanotraps for the following experiments to maximize viral capture and macrophage phagocytosis.

### Nanotraps neutralize SARS-CoV-2 infection *in vitro*

We next generated multiple types of Nanotraps to test their efficacy. First, avi-tagged biotinylated ACE2 was conjugated to the Nanotrap surface via biotin-streptavidin interactions to make Nanotrap-ACE2. In addition, we synthesized Nanotrap-Antibody by conjugating a SARS-CoV-2 neutralizing antibody to the Nanotrap surface via N-hydroxysuccinimide (NHS) esters (see details in Experimental procedures). Finally, in order to test the specificity of the Nanotraps, we made Nanotrap-Blank without a virus-binding epitope.

We next examined whether the Nantraps could effectively capture and contain SARS-CoV-2 *in vitro*. All three Nanotraps were incubated with SARS-CoV-2 spike pseudotyped lentivirus or vesicular stomatitis virus (VSV) for 1 hour before adding to HEK293T-ACE2 cells for 24 hours and 72 hours, respectively. Both Nanotrap-ACE2 and Nanotrap-Antibody completely blocked SARS-CoV-2 pseudovirus infection, while the Nanotrap-Blank did not, indicating both the specificity and functionality of our Nanotraps (Figure 3A-B and Figure S3A-B). In sharp contrast, soluble recombinant ACE2 protein only partially inhibited infection to HEK293T-ACE2 cells with both SARS-CoV-2 spike pseudotyped lentivirus (Figure 3C and Figure S3C) and VSV (Figure 3D and Figure S3D), despite the previous use of soluble ACE2 protein to inhibit SARS-CoV-2 infection(Lei et al., 2020; Monteil et al., 2020).

**Figure 3:**
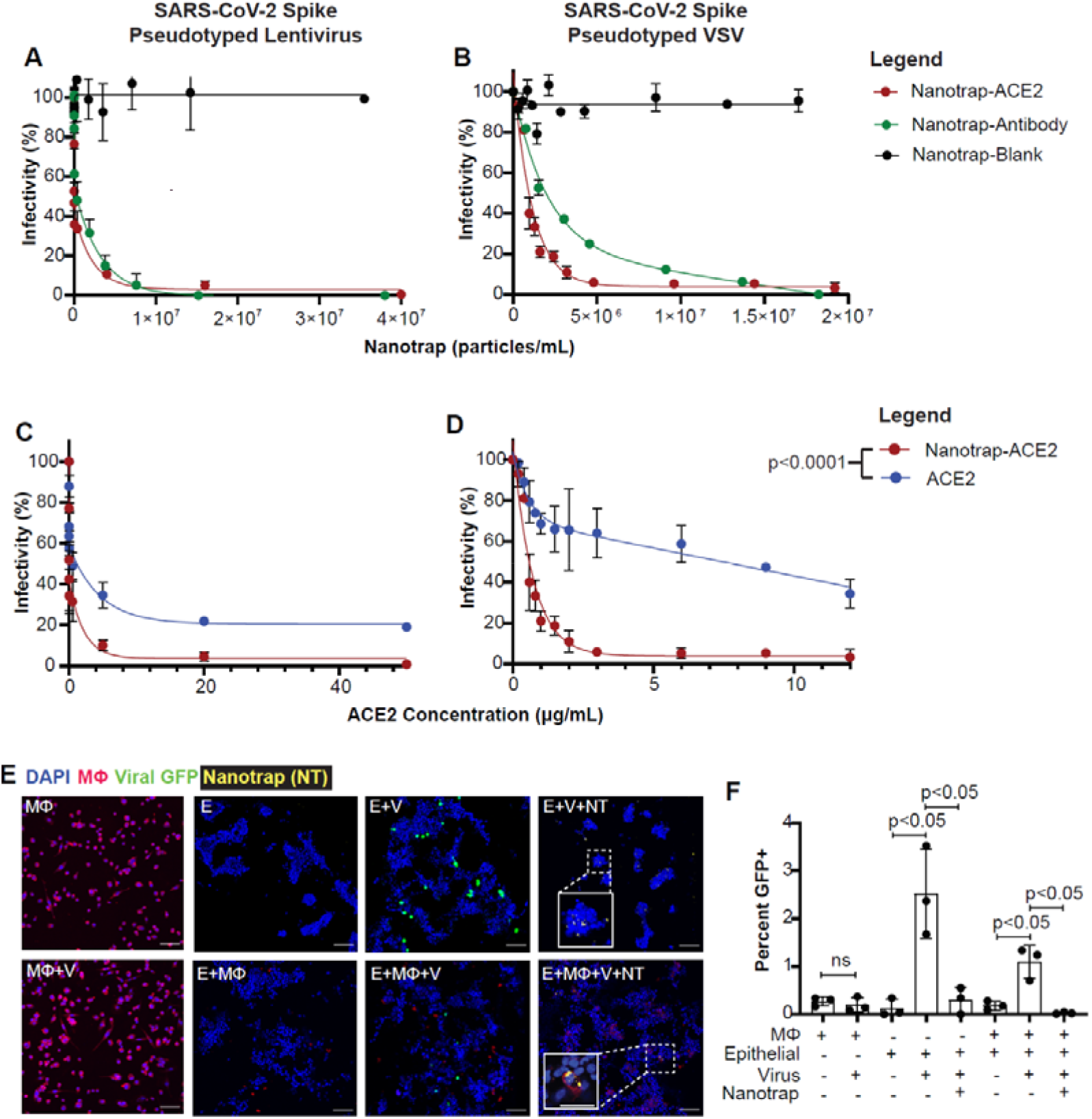
Inhibition of SARS-CoV-2 viral infection of host cells. (**A-B**) HEK293T-ACE2 cells were treated with SARS-CoV-2 spike pseudotyped lentivirus (A) or VSV (B) and Nanotrap-ACE2, Nanotrap-Antibody, or Nanotrap-Blank for 72 and 24 hours, respectively. Data are presented as mean ± SD and fitted with a two-phase decay model. (**C-D**) HEK293T-ACE2 cells were treated with SARS-CoV-2 spike pseudotyped lentivirus (C) or VSV (D) and Nanotrap-ACE2 or soluble ACE2 for 72 and 24 hours, respectively. Data are presented as mean ± SD and fitted with a trend curve. For both SARS-CoV-2 spike pseudotyped lentivirus (C) and VSV (D), the Nanotrap-ACE2 and soluble ACE2 curves differ with p<0.0001, as tested by sum-of-squares F tests. (**E**) Confocal microscopy of pseudotyped VSV infection (GFP, green) in dTHP1 macrophages (WGA, red) and A549 epithelial cells (DAPI, blue); Nanotrap-ACE2 displayed in yellow. Scale bars represent 100 μm, with inset scale bars representing 40 μm. MΦ: macrophages; V: virus; E: epithelial cells; NT: Nanotraps. (**F**) Quantification of (E). Data are shown as mean ± SD; unpaired t tests were conducted from three independent experiments.

Macrophages play a key role in controlling SARS-CoV-2 infection(Lund et al., 2016). We thus further determined whether human macrophages could efficiently engulf and degrade the virus-bound Nanotraps-ACE2 without becoming infected (Figure 1E-F). Importantly, after incubating SARS-CoV-2 spike pseudotyped VSV with dTHP-1 macrophages for 24 hours, no infection was found in the macrophages (Figure 3E-F, comparing “MΦ” with “MΦ +V”). This experiment demonstrated the feasibility of utilizing macrophages to clear the viral infection. We then infected a human lung epithelial cell line A549, which expresses physiological levels of surface ACE2, with SARS-CoV-2 spike pseudotyped VSV in the absence or presence of dTHP-1 macrophages. These data suggest that macrophages significantly reduced the viral infection but could not completely eradicate it (Figure 3E-F, comparing “E+V” with “E+ MΦ+V”). Finally, we determined whether our engineered Nanotraps triggered phosphatidylserine-mediated phagocytosis by dTHP-1 macrophages for the clearance of virus. After adding Nanotrap-ACE2 into the co-culture of epithelial cells, macrophages, and SARS-CoV-2 spike pseudotyped VSV, the viral infection was completely inhibited (Figure 3E-F, comparing “E+V+NT” and “E+MΦ +V+NT” with “E+MΦ+V”). We further observed incorporation of Nanotrap-ACE2 into the macrophage cell body, indicating successful phagocytosis (Figure 2E, Figure 3E “E+MΦ+V+NT” inset, Supplemental Video 1).

In brief summary, our *in vitro* neutralizing experiments demonstrated that our Nanotraps not only served as a sponge to capture and contain SARS-CoV-2, but also utilized the phagocytosis and sterilization machinery of macrophages to defend the host cells from infection, as we depicted in our original experimental design (Figure 1A).

### *In vivo* local delivery to lungs and biosafety profile of Nanotraps in mice

In order to assess the safety of Nanotrap treatment, we first examined *in vitro* cytotoxicity on human cell lines. Neither A549 nor HEK293T-ACE2 cells displayed significant cytotoxicity with the addition of Nanotrap-Blank, Nanotrap-ACE2, or Nanotrap-Antibody, as evaluated by a CCK8 cytotoxicity assay (Figure S4).

We next examined the delivery of Nanotraps to mouse lungs and evaluated the biosafety of Nanotraps *in vivo*. We intratracheally injected immunocompetent mice with Nanotrap-ACE2 (labelled with DiD) at a dose of 10 mg/kg. Mice were sacrificed 3 days post-injection. Delivery of Nanotraps to mouse lungs was confirmed with cryosectioned mouse lung tissues: significant Nanotrap accumulation and distribution were found in the lung tissues, particularly in regions around bronchioles in the respiratory tracts. As expected, no Nanotraps were found in the lungs of PBS-treated mice (Figure 4A-B).

**Figure 4:**
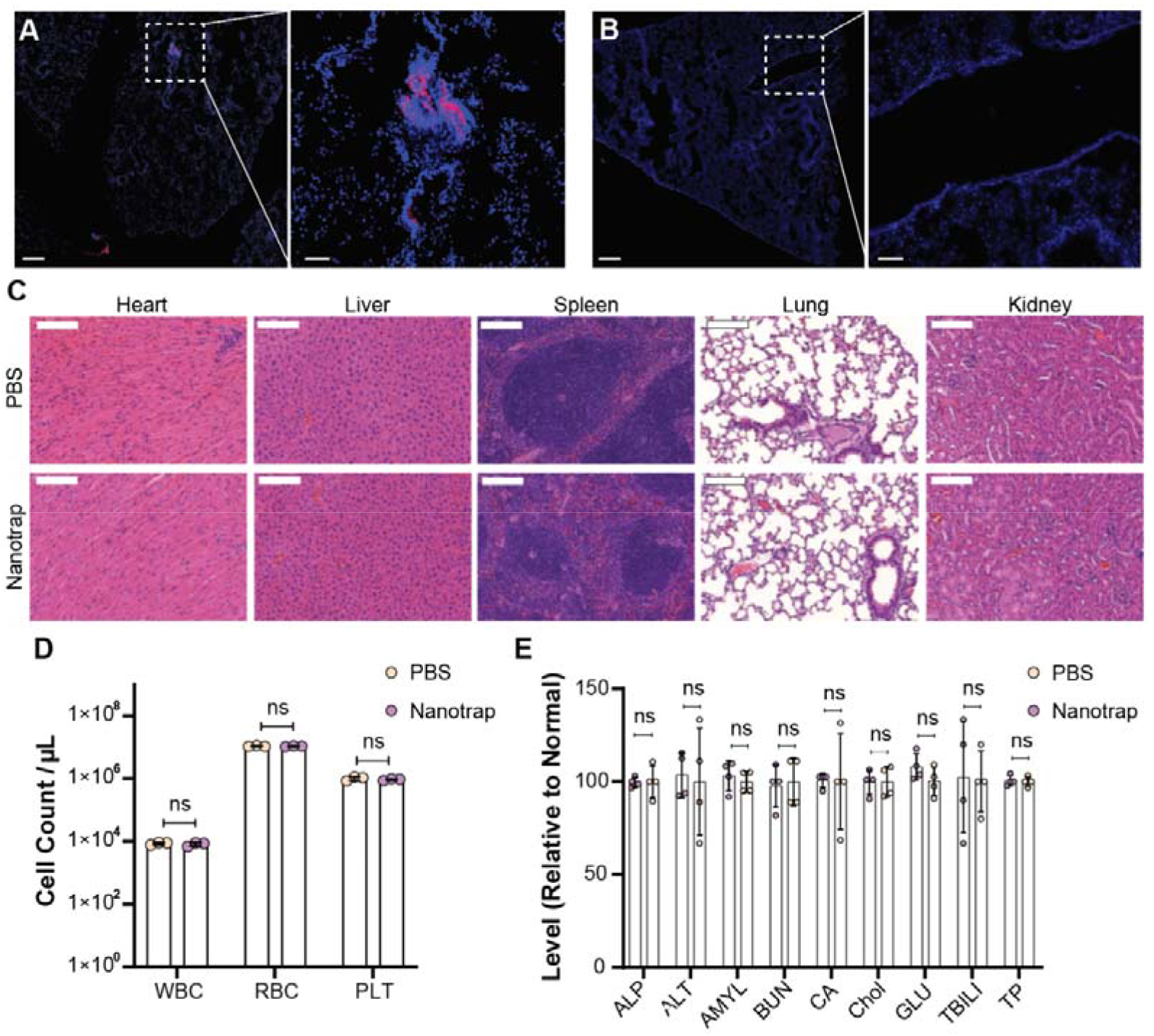
Murine *in vivo* biosafety profile of Nanotrap treatment. WT B6 mice were treated intratracheally with 10 mg/kg Nanotrap-ACE2 or PBS for 72 hours (n=4, 2 male + 2 female). (**A-B**) Representative fluorescent images of Nanotraps (DiD, red) accumulating in the lung tissues (DAPI, blue) of NT-treated mice (A) or PBS-treated mice (B) 72 hours post-intratracheal administration. Scale bars represent 250 μm; inset scale bars represent 50 μm. (**C**) Representative H&E staining of major organs sections, including heart, liver, spleen, lung and kidney. Scale bars represent 500 μm. (**D**) Blood cell counts of white blood cells (WBC), red blood cells (RBC) and platelets (PLT) 72 hours after Nanotrap or PBS treatment. Data are shown as mean ± SD; unpaired t tests were conducted from 4 replicates. (**E**) Comprehensive blood chemistry panels comparing Nanotrap- and PBS-treated mice. ALP: alkaline phosphatase; ALT: alanine aminotransferase; AMYL: amylase; BUN: urea nitrogen; CA: calcium; Chol: cholesterol; GLU: glucose; TBIL: total bilirubin; CTP: total protein.

*In* vivo safety was next analyzed. Hematoxylin and eosin (H&E) staining of major organs including lung, heart, liver, spleen, and kidney showed no histological differences in the Nanotrap-treated mice when compared to the PBS-treated control group (Figure 4C). Furthermore, complete blood counts were performed to evaluate white blood cells (WBCs), red blood cells (RBCs) and platelets (PLTs). The cell counts were similar between Nanotrap- and PBS-treated groups (Figure 4D). Next, comprehensive metabolic panels of mouse blood sera were examined to provide an overall picture of the chemical balance and metabolism. No statistical differences were found between Nanotrap- and PBS-treated mice for glucose levels, electrolyte and fluid balance, kidney function, or liver function (Figure 4E). These results demonstrated the safety of Nanotraps when delivered *in vivo*.

### The therapeutic efficacy of Nanotraps in *ex vivo* human lungs

We next examined their therapeutic efficacy in inhibiting pseudotyped SARS-CoV-2 infection in healthy, non-transplantable human donor lungs using an *ex vivo* lung perfusion (EVLP) system (Figure 5A and Supplemental Video 2). EVLP allows a lung to be perfused and ventilated *ex vivo* after organ retrieval by maintaining lungs at normothermic physiologic conditions and is thus an excellent platform to model lung diseases(Doane et al., 2012).

**Figure 5:**
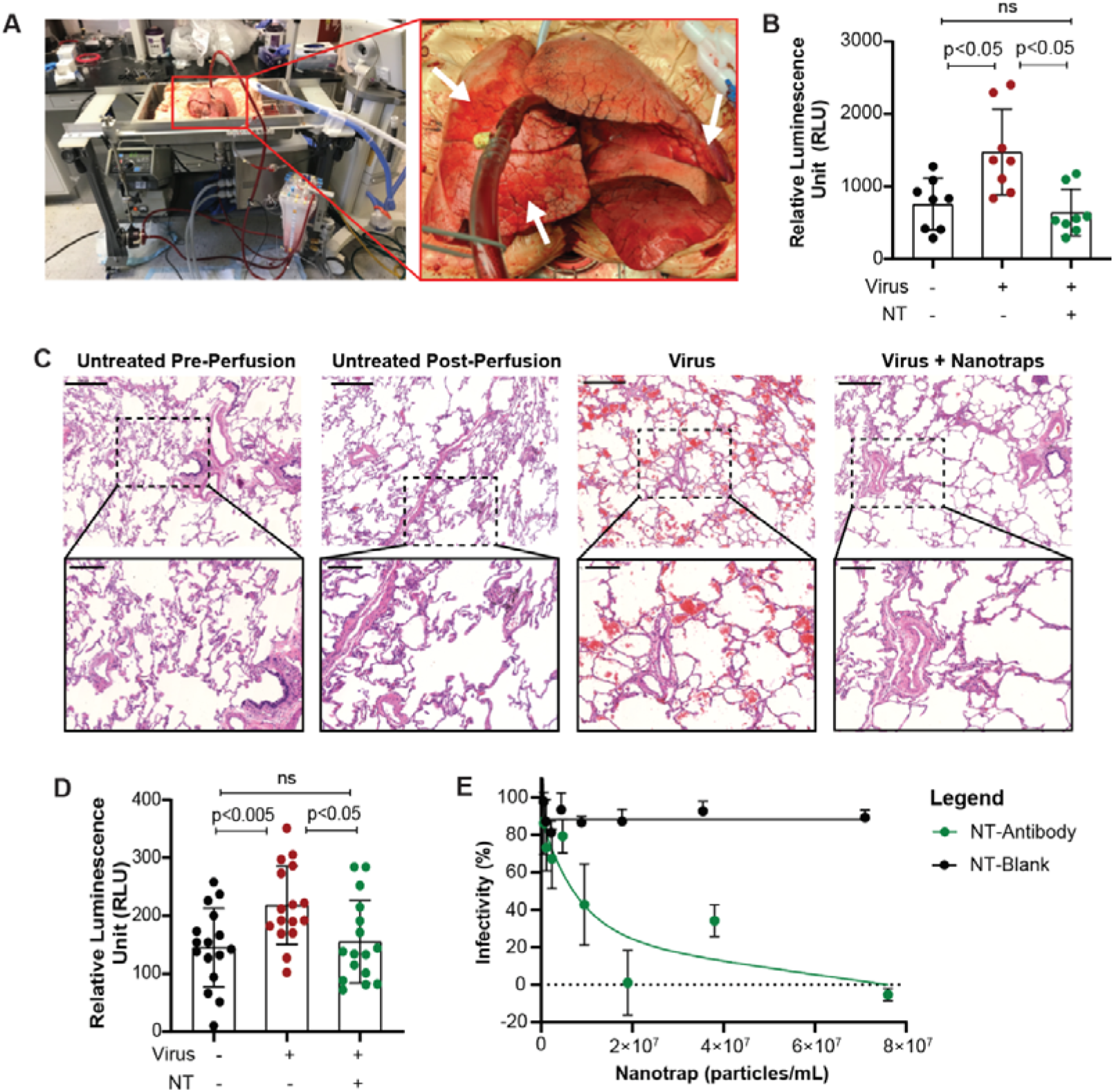
*Ex vivo* human lung perfusion system for evaluating the neutralizing ability of the Nanotrap-Antibody. (**A**) Image of human *ex vivo* lung perfusion (EVLP). Arrows indicate untreated (right upper lung), virus only (right middle lung), and virus + Nanotrap (lingula) regions. (**B**) Quantification of luciferase expression in EVLP samples 8 hours post-infection in three regions indicated in the right panel of a. Data are shown as mean ± SD. (**C**) H&E staining of EVLP. Scale bars represent 500 μm; inset scale bars represent 200 μm. (**D**) Quantification of luciferase expression in *in vitro* primary human cells 48 hours post-infection. Data are shown as mean ± SD. (**E**) Vero E6 cells treated with authentic SARS-CoV-2 and Nanotrap-Blank or Nanotrap-Antibody. Data are shown as mean ± SD fitted with a trend curve.

The infection potential of the SARS-CoV-2 spike pseudotyped lentivirus at different doses over time in primary human lung cells was first tested *in vitro*, and infection was observed within 8 hours (Figure S5A-B). After confirming infection potential, we tested our Nanotraps on an EVLP system with a pair of healthy lungs. Static lung compliance and oxygenation capacity was measured over time (Figure S5C). SARS-CoV-2 pseudovirus carrying a luciferase reporter gene was injected into the lingula of left upper lung lobe, and pseudovirus plus Nanotrap-Antibody was injected into the right middle lobe; the right upper lung lobe was used as an untreated control (Figure 5C, arrows). Human lung tissue samples were collected after perfusing for 8 hours. Single-cell suspensions were generated, and luciferase expression was determined (Figure 5B). The results showed that (1) the pseudovirus infected the lung tissues and (2) the Nanotraps completely inhibited the viral infection. Furthermore, H&E staining showed significant RBC infiltration in the virus-treated sample, which was not present in the virus plus Nanotrap-treated region (Figure 5C).

As our EVLP system maintains lung viability for less than 12 hours, we treated single-cell suspensions of healthy, untreated lung from the right upper lobe *in vitro* for 48 hours to confirm the Nanotraps can function for longer term incubations in human tissue. Again, Nanotrap-Antibody was able to fully inhibit the virus (Figure 5D). Finally, since the EVLP could not be conducted under BSL-3 conditions in order to use authentic SARS-CoV-2, we tested the ability of the Nanotraps to prevent authentic SARS-CoV-2 from infecting Vero E6 cells, which are highly susceptible to SARS-CoV-2 infection(Matsuyama et al., 2020). Indeed, Nanotrap-Antibody was able to completely inhibit infection of authentic SARS-CoV-2, as expected (Figure 5E).

Taken together, our EVLP experiments demonstrated that (1) SARS-CoV-2 pseudovirus can infect human lung, and (2) our newly engineered Nanotraps can completely block the viral infection, thus paving the way for future clinical trials using Nanotraps for the inhibition of SARS-CoV-2 infection.

## Discussion

The highly contagious SARS-CoV-2 has caused the global COVID-19 pandemic, so effective and safe treatments are urgently needed. Remdesivir has been approved by the FDA to treat severe COVID-19(Beigel et al., 2020), despite its inconsistent clinical benefits and various reported adverse effects(Grein et al., 2020; Wang et al., 2020b). Transfusion of convalescent plasma from recovered patients has shown clinical benefits in some COVID-19 patients(Shen et al., 2020), however, this approach is challenged by the limited availability of donor plasma and appropriate medical facilities(Pérez-Cameo and Marín-Lahoz, 2020). Simultaneously, tremendous efforts have been devoted to the development of vaccines, neutralizing antibodies, and other drugs for the prevention and treatment of COVID-19. While recently developed vaccines are being dispersed to the population, safe and effective medicines to treat SARS-CoV-2 infection are largely lacking. For example, Nanosponges have been developed (Zhang et al., 2020), but they cannot completely inhibit infection. They are also not a viable treatment option: as each patient’s human leukocyte antigen is different, Nanosponges would have to be personalized for each patient. Large amounts of uninfected primary cells would have to be collected from each patient to make personalized Nanosponges, similar to CAR-T therapy, which has resulted in exorbitant costs and is thus unattainable by many patients(Fiorenza et al., 2020). In addition, recombinant soluble ACE2 or ACE2-IgG proteins have been proposed for use as treatments (Chen et al., 2020b; Lei et al., 2020), but soluble proteins are known to be much more liable to degradation than nanoparticles (Yu et al., 2016). Here we aimed to develop Nanotraps to effectively inhibit SARS-CoV-2 infection. The design of our Nanotraps was inspired by the ability of tumor cells to secrete PD-L1 exosomes, which bind to and suppress T-cell immune functions and thus prevent the killing and clearance of the tumor cells(Chen et al., 2018; Daassi et al., 2020). In a similar fashion, the synthetic Nanotraps can mimic the target cells to ensnare the virus. Meanwhile, the synthetic polymer-lipid complexes takes advantage of both the stability from polymers and surface flexibility of lipids(Mukherjee et al., 2019), thereby providing a well-controlled nanomaterial with a high capacity to trap the virus by mimicking target cells. We thus created Nanotraps to bind and inhibit SARS-CoV-2 infection to host cells.

To block the interaction between the SARS-CoV-2 spike protein and the host ACE2 receptors, we coated the Nanotrap surfaces with a high molecular density of either recombinant ACE2 proteins or anti-SARS-CoV-2 neutralizing antibodies (Figure 1A). In principle, the high binding avidity, high diffusivity, and small size of Nanotraps should enable them to easily outcompete low ACE2-expressing host cells in capturing the SARS-CoV-2, thus effectively containing the viruses on their surfaces. Indeed, our experiments demonstrated that viral infection of both pseudotyped and authentic SARS-CoV-2 across human cell lines, lung primary cells and lung organs can be completely inhibited by Nanotrap-ACE2 or Nanotrap-Antibody (Figure 3 and Figure 5). Notably, Nanotrap-ACE2 was superior to the soluble recombinant ACE2 proteins in containing SARS-CoV-2(Lei et al., 2020; Monteil et al., 2020), attributing to their high binding avidity (Figure 3C-D).

Furthermore, our Nanotraps harness the immune system to clear the SARS-CoV-2 (Figure 2 and Figure 3). By incorporating the phagocyte-specific phosphatidylserine ligands onto the Nanotrap surfaces, macrophages readily engulfed the virus-bound Nanotraps without becoming infected themselves (Figure 3E-F). While macrophages were used as a proof-of-principle in this study, other professional phagocytes such as neutrophils, monocytes, and dendritic cells should be able to similarly clear the virus-bound Nanotraps. In particular, macrophages and dendritic cells are professional antigen presenting cells, which present engulfed antigens to the adaptive immune system(Janeway, 2008). Since the Nanotraps are able to engage antigen presenting cells, it is possible that they may also elicit virus-specific adaptive immune responses. Future studies will evaluate whether Nanotraps can prime adaptive immune responses, thereby promoting vaccine-like protection(Klichinsky et al., 2020).

In addition, we purposely designed Nanotraps to be biocompatible, biodegradable and safe. The Nanotraps were composed of FDA-approved polymers and lipids, which provides the possibility for safe administration in a clinical setting. Indeed, our biosafety experiments have demonstrated excellent safety profile *in vitro* and *in vivo* (Figure 4).

Lastly, we tested the efficacy of the Nanotraps in a human EVLP system. Superior to lung organoids which cannot reproduce whole-organ response to viral infection(Han et al., 2020; Monteil et al., 2020; Suzuki et al., 2020), and non-human primate models which are extremely costly(Chandrashekar et al., 2020; Rockx et al., 2020; Yu et al., 2020), the EVLP system is a clinically relevant model. We showed that our Nanotraps can completely inhibit viral infection in living human lungs (Figure 5). As current biosafety regulations preclude the testing of authentic SARS-CoV-2 in the EVLP, we further confirmed our Nanotraps can inhibit authentic virus *in vitro* (Figure 5E). These experiments together suggest our Nanotraps could potentially be used to treat SARS-CoV-2 infection in the clinic.

In summary, we developed a new type of potent, effective nanomedicine “Nanotraps” to contain and clear SARS-CoV-2 by harnessing and integrating the power of nanotechnology and immunology. The Nanotraps completely inhibited the SARS-CoV-2 infection to human cells and lung organs. The Nanotraps are effective, biocompatible, safe, stable, feasible for mass production. It is reasonable to hypothesize that the Nanotraps could be easily formulated into a nasal spray or inhaler for easy administration and direct delivery to the respiratory system, or as an oral or ocular liquid, or subcutaneous, intramuscular or intravenous injection to target different sites of SARS-CoV-2 exposure, thus offering flexibility in administration. Furthermore, the design of our Nanotrap is highly versatile: they can be modified to incorporate small molecule drugs or protein/mRNA vaccines to their core, and different human ACE2 recombinant proteins(Lei et al., 2020; Monteil et al., 2020), human anti-SARS-CoV-2 neutralizing antibodies(Cao et al., 2020; Chen et al., 2020b; Chi et al., 2020; Rogers et al., 2020; Shi et al., 2020), or any newly-developed therapeutic proteins or peptides can be conjugated to the surface, thus easily extending their applications beyond our current study. Overall, we thus expect continuous development of this nanomedicine for clinical use of preventing and treating SARS-CoV-2 infection.

## EXPERIMENTAL PROCEDURES

### Resource availability

#### Lead contact

Further information and requests for resources and reagents should be directed to and will be fulfilled by the Lead Contact, Jun Huang.

#### Materials Availability

The materials generated in this study are available from the corresponding author upon request.

#### Data and Code Availability

The data used to support the findings of this study are available from the corresponding author upon request.

### Synthesis of Nanotraps

To synthesize the Nanotraps, we employed a two-step method developed for polymer-lipid hybrid nanoparticles, in which the polymer and lipid components were prepared separately and combined at the end of the process. The PLA nanoparticles were prepared in accordance with existing methods(Li et al., 2020) through an oil-in-water emulsion solvent evaporation process. 100 mg of PLA and 100 μL of PFOB were dissolved in 3.5 mL of dichloromethane. The organic phase was mixed with 20 mL of 2.0% PVA solution. The mixture was emulsified by sonication (Fisher Sonics) on ice for 2 minutes. The dichloromethane in the emulsified mixture was evaporated under magnetic stirring at 300 rpm for 3-4 hours at room temperature. The resulting solution was centrifuged and the pellets were washed with PBS 3 times (5,000×g for 10 minutes), and then the pellets were lyophilized and stored at 4 °C before use. For any fluorescence labeling, 0.05 mg DiD or DiO was added into the 100 mg PLA organic phase when preparing the Nanotraps. For PLA nanoparticles with size 200 nm, 100 mg of PLA was emulsified with 2.0% PVA and the supernatant was collected after centrifugation. For PLA nanoparticles with size 500 nm, 100 mg of PLA and 100 μL PFOB were mixed, then emulsified with 2.0% PVA and the pellets were collected after centrifugation. For PLA nanoparticles with size 1200 nm, 100 mg of PLA and 100 μL PFOB were mixed and emulsified with 1.0% PVA and the pellets were collected after centrifugation.

Functionalized PLA-nanoparticles composed of 15% phosphatidylserine and 0.5% DSPE-PEG2000-biotin were prepared by dissolving DOPS (0.388 μmol), DSPC (1.292 μmol), cholesterol (0.775 μmol), DSPE-mPEG_2000_ (0.166 μmol) and DSPE-PEG2000-biotin (0.013 μmol) at a molar ratio of 10:120:60:9:1 in dichloromethane. The dichloromethane solvent was slowly evaporated by heating the lipid solution at 55 °C to remove the solvent and further dried in a vacuum drying oven to produce a dried lipid film. The lipid film was reconstituted in 2 mL of PBS (pH 7.4) containing 0.2 mg PLA nanoparticles and the contents were hydrated at 60 °C under ultrasonication (Branson CPX5800H). Then the mixture was sonicated with a sonicator probe (Fisher Sonics) for 2 minutes (100 W, 22.5 kHz, 30% amplitude). Formulations for Nanotraps with different phosphatidylserine surface densities were listed in Table S1.

To synthesize Nanotrap-ACE2, 1 mg of the biotin functionalized PLA nanoparticles were then incubated with streptavidin (34.25 μL, 2 g/L) on ice for 40 minutes under magnetic stirring. The resulting solution was centrifuged at 5,000×g for 10 minutes and washed with 1 mL PBS for 3 times to remove excess streptavidin. The pellet was resuspended in 100 μL PBS and incubated with biotinylated ACE2 (Bioss Antibodies) for 30 minutes on ice. The resulting PLA@DOPS/ biotin~SA~ACE2 nanoparticle was termed as Nanotrap-ACE2.

Nanotrap-Antibody was synthesized by combining 15% DOPS (0.313 mg, 0.386 μmol), 50% DSPC (1.018 mg, 1.287 μmol), 30% cholesterol (0.300 mg, 0.773 μmol) and 5% DSPE-PEG_2000_-NHS (0.370 mg, 0.129 μmol) in dichloromethane. The lipids mixtures were vacuum dried overnight, and the resulting thin film was hydrated with PLA-NPs PBS solution under ultrasonic water bath and further reacted with 100 μg of SARS-COV-2 neutralizing antibody for 4 hours at 4 °C.

### Characterization of Nanotraps

The sizes of Nanotraps were measured by a dynamic light scattering particle size analyzer (Malvern Zetasizer). Briefly, 1 μL of Nanotraps were dispersed in 0.1× PBS and further dispersed in ultrasonic water bath for 10 minutes before testing. The size measurement was carried at 25 °C with count rates within 300-500 kcps and measured 3 times. The zeta potentials of Nanotraps were performed by a Möbiuζ system (Wyatt Technology). The data were presented as mean ± SD.

AF488-labeled anti-ACE2 antibody (Santa Cruz Biotechnology) was added to the Nanotrap-ACE2 for 30 minutes on ice and centrifuged at 5,000×g for 10 min, the pellet was washed with PBS for 3 times. The resulting Nanotraps were resuspended in 50% glycol and imaged by total internal reflection fluorescence microscopy (Nikon) with 488 nm and 647 nm excitation lasers and 200 milliseconds exposure. Line scans were performed in Fiji.

The sizes and morphologies of the Nanotraps were studied by scanning electron microscopy. Briefly, 10 μL of Nanotrap-ACE2 were diluted in MiliQ water and further dispersed on ultrasonic water bath for 10 minutes before adding onto a silicon chip. 40 μL of SARS-CoV-2 spike pseudotyped lentivirus was fixed by 4% PFA at 37 °C for 30 minutes, the virus was then washed by PBS 3 times using an Amicon Ultra-15 Centrifugal Filter (pore size 100 KDa) at 3,000×g, 10 minutes. The resulting fixed virus was incubated with 10 μL Nanotraps-ACE2 at 37 °C for 1 hour and added onto the 1 cm^2^ silicon chip followed by airdrying overnight. After dehydration, the samples were coated with 8 nm platinum/palladium by sputter coater (Cressington 208HR). The scanning electronic microscope (Carl Zeiss Merlin) was used to image the morphology of the Nanotraps with an accelerating voltage of 2.0 kV. For each sample, more than 10 measurements with different magnification were performed to ensure the repeatability of the results.

### Flow cytometry analysis of Nanotraps phagocytosis

Macrophage differentiation was conducted as follows(Hsu et al., 1996): THP-1 cells were treated with 150 nM PMA for 24 hours and replaced by fresh culture medium for another 24 hours. The differentiated THP-1 cells were harvested as dTHP-1 macrophages and maintained in RPMI supplemented with 10% FBS and 1% Penicillin-Streptomycin. The dTHP-1 macrophages were then released from the plate with Trypsin-EDTA (0.25%) and cell number was counted by a hemocytometer. 1 million dTHP-1 macrophages were seeded into each 6-well plate overnight, and DiO-labeled Nanotraps (200, 500, and 1200 nm) were incubated with dTHP-1 macrophages for 0, 24, or 48 hours. For phosphatidylserine investigation, dTHP-1 macrophages were incubated with Nanotraps containing different phosphatidylserine molar ratios (0%, 5%, 10%, 15%) for 0, 2, 4, 6, 24, or 48 hours. The cells were harvested and washed 3 times with PBS at 300×g for 5 minutes, and stained with Zombie NIR™ Fixable Viability Kit (BioLegend) on ice for 10 minutes. Then cells were washed with FACS buffer (PBS, 10% FBS, 0.1% NaN_3_) 2 times and resuspended in 200 μL FACS buffer. Flow cytometry was carried on a BD LSRFortessa™ Flow Cytometer. Live and single cells were gated and DiO fluorescence channel was used to indicate the phagocytosis efficiency of different Nanotraps. The data were further analyzed by FlowJo (BD) and Prism (Graphpad) softwares.

### Lattice light-sheet microcopy imaging analysis of Nanotrap phagocytosis

For lattice light-sheet imaging, 4×10^4^ dTHP-1 macrophages were seeded onto each coverslip and DiD-labeled Nanotraps with different phosphatidylserine molar ratios (0%, 10%, and 15%) were added for 24- or 48-hour incubation. The cells were washed by PBS for 3 times, fixed with 4% PFA, stained with (5 μg/mL) CF488 Wheat Germ Agglutinin (WGA) Conjugates (Biotium), and washed 3 times with HBSS. Coverslips were imaged by lattice light-sheet microscopy (3i) using z+objective scanning. Imaging was conducted with 488 nm and 647 nm lasers, with dither set to 3 and 20 millisecond exposures. Z-steps (60) were collected with a 0.4 μm step size. Resulting images were deconvolved as described previously(Rosenberg and Huang, 2020; Rosenberg et al., 2020) using LLSpy (cudaDeconv) used under license from Howard Hughes Medical Institute, Janelia Research Campus. Image reconstruction videos were made in Imaris (Bitplane).

### Production of SARS-CoV-2 spike pseudotyped VSV

Packaging cells (HEK293T) in serum-free DMEM were transfected with 9 μg pCAGGS SARS-CoV-2 spike expression plasmid using polyethylenimine (PEI). After 24 hours, 3×10^7^ FFU VSVdG*G-GFP virus was added to the HEK293T cells and incubated for another 48 hours. Media were collected and spun at 500×g for 3 minutes to remove cell debris, then passed through a 0.45 μm pore filter. The virus was then stored in −80 °C. To check the infection rate, HEK293T-ACE2 cells (Integral Cat# C-HA102) were incubated with the final VSVdG-GFP*CoV2 pseudovirus and visualized by the presence of GFP positive cells through direct microscopic imaging or flow cytometry.

### SARS-CoV-2 pseudovirus neutralizing assay

For SARS-CoV-2 spike pseudotyped VSV neutralizing assay, 4×10^4^ HEK293T-ACE2 cells (maintained in DMEM supplemented with 10% FBS and 1% Penicillin-Streptomycin) were seeded in a 96-well plate overnight. Different concentrations of ACE2 proteins or Nanotraps (containing 0.1, 0.2, 0.4, 0.6, 0.8, 1.0, 1.5, 3.0, 6.0, 9.0, 12, 15 μg/mL ACE2) were incubated with 500 FFU SARS-CoV-2 spike pseudotyped VSV for 1 hour in 37 °C. The virus-Nanotrap solution was added into the HEK293T-ACE2 cells and incubated for 24 hours (n=3 for each group). The cells were imaged with a fluorescence microscope (Nikon) using 10×/0.30 NA objective. The excitation wavelengths were 470 ± 25 nm (Spectra X, Lumencor). The emissions of GFP were captured by an Andor iXon Ultra 888 back-illuminated EMCCD camera (Oxford Instruments). The number of GFP-positive cells were counted manually by objective three times. The viral infection rates were calculated as the ratio of GFP-positive cells in the group incubated to that of the group incubated with virus alone.

For SARS-CoV-2 spike pseudotyped lentivirus neutralizing assay, 1×10^4^ HEK293T-ACE2 cells were seeded onto a 384-well plate overnight. Nanotraps or ACE2 was added to SARS-CoV-2 pseudovirus (4 μL per well) and incubated for 1 hour at 37 °C. The virus-Nanotrap solution was added to each well (n = 3 for each group). 72 hours later, the plate was centrifuged for 5 minutes at 500×g to prevent cell loss. Supernatant was aspirated and 35 μL of PBS was added. PBS was carefully aspirated, leaving ~15 μL of liquid behind. Renilla-Glo assay substrate was added to the Assay Buffer at a 1:100 dilution, then 15 μL of the substrate:buffer was added to each well of a 384-well plate. The bioluminescence was recorded by a microplate reader (Fisher Scientific BioTek Cytation 5) with an exposure of 200 milliseconds. Wells infected with pseudovirus only were normalized as 100%.

### Co-culture assay

THP-1 cells were differentiated into macrophages as described above. Coculture was carried out in a macrophage to A549 cell ratio of 1:5. 4×10^4^ A549 cells were seeded in an 18-well microslide (Vivid) overnight and 8×10^3^ dTHP-1 macrophages were added onto the A549 cells for another 6 hours. 500 FFU SARS-CoV-2 spike pseudotyped VSV was incubated with Nanotraps or PBS in 37 °C for 1 hour before adding to the coculture cells. 24 hours later, the cells were fixed with 4% PFA and stained with CF532 Wheat Germ Agglutinin (WGA) Conjugates (Biotium) and DAPI and imaged under a confocal microscope (Leica SP8). Percent infectivity was quantified in FIJI by dividing GFP^+^ cells by total cell number (DAPI-stained nuclei). Each channel was processed as follows: Image>Threshold (“Huang” preset(Huang and Wang)); Image>Binary>Fill Holes; Image>Binary>Watershed; Analyze>Count Particles.

### *In vitro* cytotoxicity assay

A549 or HEK293T-ACE2 cells (both maintained in DMEM supplemented with 10% FBS and 1% Penicillin-Streptomycin) were seed in a 96-well plate at a density of 1×10^4^ cells/well in 100 μL of culture medium overnight. Nanotraps (3.8×10^7^ particles/mL) were added into cells and the cells were cultured in a CO_2_ incubator at 37 °C for 72 hours.10 μL of CCK-8 (MedChem Express) solution was added to each well of the plate. The plate was incubated for 2 hours in the incubator. Then it was put into a microplate reader (Fisher Scientific BioTek Cytation 5) and the plate was gently shaken for 1 minute before measuring the absorbance at 450 nm. The cytotoxicity was calculated by cell viability, that the relative absorbance from the control wells without Nanotraps were normalized as 100%.

### *In vivo* biosafety assays

C57BL/6NHsd mice at the age of 6 weeks were purchased from Envigo and maintained at the Animal Facility of the University of Chicago. The animal study protocols were approved by the Institutional Animal Care and Use Committee (IACUC) of University of Chicago. To evaluate the safety of the Nanotraps, 2 male and 2 female 6-10-week-old C57BL/6NHsd mice were intratracheally administered with 10 mg/kg Nanotrap-ACE2 in 50 μL PBS. Blood samples were collected by submandibular vein via cheek punch using a commercially available 4 mm point lancet after three days. A small aliquot of approximately 100 μL blood was collected into EDTA-containing heparinized tubes, and red blood cells, white blood cells, and platelets were counted by a Hematology Analyzer (Beckman Coulter Act Diff 5 CP) according to the manufacturer’s instructions. For the comprehensive chemistry panels, blood was allowed to coagulate in 4 °C for 2 hours, and serum was collected after centrifugation (1,000×g for 15 minutes) for analysis. Serum alkaline phosphatase, alanine aminotransferase, amylase, urea nitrogen, calcium, cholesterol, glucose, total bilirubin, total proteins were determined by a Vet Axcel blood chemistry analyzer (Alfa Wasserman). Lungs, heart, liver, spleen, and kidney were collected from the same mice, fixed in 10% formalin for 24 hours, embedded in paraffin. The resulting blocks were cut in 5 μm sections and further stained with hematoxylin and eosin (H&E) by the University of Chicago Human Tissue Research Center. Fluorescent imaging samples were collected in Tissue-Tek® O.C.T. Compound on dried ice and stored in −80 °C before cryosectioning on a cryostat (Leica). The obtained 10 μm thick tissue slides were then further stained with DAPI. The histology and fluorescence slides were scanned by a CRi Pannoramic MIDI 20× whole slide scanner and analyzed on the QuPath software.

### *Ex vivo* lung perfusion assay

Non-transplantable human lungs were obtained from deceased individuals provided by the organ procurement organization Gift of Hope. All specimens and data were de-identified prior to receipt. This study was deemed exempt by the University of Chicago Institutional Review Board (IRB19-1942).

#### Lung Harvest

Living lungs unsuitable for transplantation were harvested in standard clinical fashion(Hsu et al., 1996) from deceased patients. Figure 5A shows the lung of a 56-year-old male patient (87.2 kg, cause of death: brain death). Lungs were transported to the laboratory at 4 °C.

#### Lung inoculation

Tissue samples were collected from the edge of the right superior lobe before perfusion as a untreated control; 500 μL SARS-CoV-2 spike pseudotyped lentivirus (Intergral Molecular, RVP 701) was resuspended in 5 mL PBS and injected into the lingula of the left lung; 500 μL SARS-CoV-2 spike pseudotyped lentivirus was first incubated with Nanotrap-Antibody carrying 250 μg neutralizing antibody for 1 hour at 37 °C before inoculation. This mixture was then injected into the right middle lobe. Tissue samples at all three sites were collected. Some samples were immersed in MACS buffer for tissue dissociation, and other samples were fixed in 4% PFA, sliced and stained for H&E.

#### Lung perfusion

The lung bloc was perfused according to published techniques(Cypel et al., 2008; Ross et al., 2019). A centrifugal pump was used to perfuse the pulmonary artery with deoxygenated cellular perfusate (1×Dulbecco’s Modified Eagle Medium containing 4.5g/K D-glucose, L-glutamine and 110 mg/L of sodium pyruvate, with addition of 5% bovine serum albumin and 2 units of packed red blood cells). The left atrium was left open for gravity drainage. The trachea was incubated and the lung was ventilated with volume control ventilation (tidal volume set at 6-8 mL/weight of ideal body weight (kg) of donor, respiratory rate of 8-13, and fraction of inspired oxygen set at 21%). Sweep gas composed of 8% CO_2_, 3% O_2_, 89% N_2_ was connected to the hollow-fiber de-oxygenator heat exchanger to remove oxygen and add in CO_2_ into the perfusate returning back to the lung. After initiation of perfusion with gradual warming and increasing pump flow over 30 minutes, the lung bloc was maintained at 37 °C with pump flow calibrated to pulmonary artery pressure of 10 – 20 mmHg for 8 hours.

#### Sampling

Perfusate was sampled at serial time points from the pulmonary artery and left atrium. Differences in oxygen content between perfusate samples were used to calculate the oxygenation capacity of the lung. Airway pressure was measured periodically to calculate lung compliance based on tidal volume. Tissue samples were collected from treated and untreated lobes of the lung at time 0 and time 8 hours of perfusion.

#### Sample Processing

After 8 hours perfusion, tissues were harvested as described above. Tissue dissociation was performed by mechanical digestion in DMEM media treated with 2.5 U/mL DNase (company); samples were passed through 70 μm cell strainers (Fisher Scientific). Red blood cells were lysed in 10 mL RBC lysis buffer (Life Technologies), washed 3 times in DMEM (300×g for 3 minutes), and resuspended in 10 mL DMEM. Cells were counted with a hemocytometer and immediately used for luciferase assay or cryopreserved in Cell Banker medium (Amsbio) at a density of 4×10^6^ cells/mL.

For luciferase assay, the cells harvested as described above were washed with PBS (300×g for 3 minutes) 3 times. Then 2×10^4^ cells were seeded into 96-well plate. Renilla-Glo Assay Substrate was added to the Assay Buffer at a 1:100 dilution, then 30 μL of the Substrate Buffer was added to each well for 10 minutes. The bioluminescence from each well was detected by a microplate reader (Fisher Scientific BioTek Cytation 5) with an exposure of 200 milliseconds.

Primary cells harvested from the untreated right superior lobe of the human lung were further used for infection analyses. Briefly, 1×10^4^ primary lung cells were seeded onto a 384-well plate. 20 μL Nanotraps-Antibody (3.8×10^7^ particles/mL) was added to 10 μL SARS-CoV-2 spike pseudotyped lentivirus (Intergral Molecular, RVP 701) per well and incubated for 1 hour at 37 °C before adding to the cells. For virus only group, 10 μL SARS-CoV-2 spike pseudotyped lentivirus was added to the cells. 48 hours later, the plate was centrifuged for 5 minutes at 500×g to prevent cell loss. Supernatant was aspirated and 35 μL PBS was added. PBS was carefully aspirated, leaving ~15 μL of liquid behind. Renilla-Glo Assay Substrate was added to the Assay Buffer at a 1:100 dilution, then 15 μL of the Substrate:Buffer was added to each well of a 384-well plate. The bioluminescence was recorded by a microplate reader (Fisher Scientific BioTek Cytation 5) with an exposure of 200 milliseconds. The infectivity was calculated by the relative luminescence intensity: wells infected with SARS-CoV-2 pseudovirus only were normalized as 100%.

### Authentic SARS-CoV-2 Neutralizing Assay

All SARS-CoV-2 infections were performed in biosafety level 3 conditions at the University of Chicago Howard T. Ricketts Regional Biocontainment Laboratory. African green monkey kidney (Vero E6) cells were maintained in DMEM supplemented with 10% FBS and 1% Penicillin-Streptomycin. Nanotraps or neutralizing antibodies were serially diluted 2-fold and mixed with 400 PFU of SARS-CoV-2 (nCoV/Washington/1/2020, kindly provided by the National Biocontainment Laboratory, Galveston, TX) for one hour at 37 ºC, then used to infect Vero E6 cells for three days. Cells were fixed with 3.7% formalin and stained with 0.25% crystal violet. Crystal violet stained cells were then quantified by absorbance at (595 nm) with a Tecan m200 microplate reader. Cell survival was calculated by normalizing untreated cells to 100%.

## SUPPLEMENTAL INFORMATION

Supplemental information can be found online at https://doi.org/xxxxxxxxxx

## ACKNOWLEDGEMENTS

This work was supported by NIH New Innovator award 1DP2AI144245 (J.H.), NSF Career award 1653782 (J.H.) and NIDDK RC2DK122394 (E.C.). M.C. is partially supported by the UChicago Big Ideas Generator Grant. J.R. is supported by the NSF Graduate Research Fellowships Program DGE-1746045. We would like to thank Dr. Nicholas Ankenbruck, Yifei Hui, Guoshuai Cao, Ruyi Wang, Joe Reda and Aaron Alpar for their experimental guidance. P.P.M. is supported by DP2DA051912. We would also like to thank Dr. Aaron Esser-Kahn for kindly providing THP-1 cells and Dr. Jeffery Hubbell for the use of laboratory equipment. We also thank the support from the University of Chicago Human Tissue Research Center, University of Chicago Integrated Light Microscopy Core, and the Soft Matter Characterization Facility of the Pritzker School of Molecular Engineering for sample processing and characterization.

## AUTHOR CONTRIBUTIONS

Conceptualization: M.C. with input from J.H. Methodology: M.C. with input from J.H. Investigation/experiments: M.C., J.R., X.C., A.C.H.L., J.S., M.N., T.W., V.M., A.J.E., J.F., J.S.D., K.S., G.R., B.T., and M.L.M. SARS-CoV-2 pseudovirus generation and discussions: P.P.M. Formal Analysis: M.C. and J. R. with input from J.H. Manuscript writing: M.C., J.H., and J.R.. Funding acquisition: J.H. and E.C. Supervision: J.H. Manuscript editing and review: all authors.

