## Supplementary material for "Nanotraps for the containment and clearance of SARS-CoV-2": SI

### Supplemental Methods

#### Chemicals and reagents

Polylactic acid (PLA), polyvinyl alcohol (PVA), cholesterol, 1-bromoheptadeca fluorooctane (PFOB), dichloromethane, 3,3′-dioctadecyloxacarbocyanine perchlorate (DiO), dimethyl sulfoxide(anhydrous) and phorbol 12-myristate 13-acetate (PMA) were purchased from Sigma-Aldrich. 1,2-distearoyl-sn-glycero-3-phosphocholine (DSPC), 1,2-dioleoyl-sn-glycero-3-pho spho-L-serine (sodium salt) (DOPS), 1,2-distearoyl-sn-glycero-3-phosphoethanolamine-N-[methoxy(polyethylene glycol)-2000] (DSPE-mPEG$_{2000}$), 1,2-distearoyl-sn-glycero-3-phosphoethanolamine-N-[biotinyl (polyethylene glycol)-2000] (ammonium salt) (DSPE-PEG$_{2000}$-biotin), 1,2-distearoyl-sn-glycero-3-phosphoethanolamine-N-[carboxy(polyethylene glycol)-2000, NHS ester] (DSPE-PEG$_{2000}$-NHS) were purchased from Avanti Polar Lipids. Renilla-GLO luciferase substrate was purchased from Promega. 1,1-dioctadecyl-3,3,3,3-tetramethylindodicarbocyanine (DiD), wheat germ agglutinin CF™488A conjugate and wheat germ agglutinin Cf532 conjugates were ordered from Biotium. Zombie NIR™ Fixable Viability Kit was ordered from BioLegend. Cell Counting Kit-8 (CCK8) was purchased from MedChem Express. 384-well tissue culture plates were purchased from Santa Cruz Biotechnology. Biotinylated recombinant human ACE2 was purchased from Bioss Antibodies. ACE2 (E-11) Alexa Fluor 488 was purchased from Santa Cruz Biotechnology (cat# sc-390851 AF488, RRID: AB_2861379). Mouse anti-SARS-CoV-2 (2019-nCoV) Spike Neutralizing Antibody was purchased from SinoBiological (cat#40592-MM57, RRID: AB_2857935). SARS-CoV-2 spike pseudotyped lentivirus with luciferase reporter gene was purchased from Integral Molecular (Catalog# RVP 701). Amicon Ultra-15 centrifugal filter unit with Ultracel-100 membrane (100 KDa, 4 mL) was purchase from EMD Millipore. 32% paraformaldehyde aqueous solution was purchased from Electron Microscopy Sciences.

#### Cell lines

HEK293T-ACE2 cells were provided by Integral Molecular (Integral Cat# C-HA102) and were cultured by cell culture media containing DMEM + 10% FBS + 10 mM HEPES + 1% Penicillin-Streptomycin. Human lung epithelial cell line A549 cells were obtained from (ATCC® CCL-185™) and cultured in DMEM + 10% FBS + 1% Penicillin-Streptomycin. THP-1 cells were kindly provided by Dr. Aaron Esser-Kahn and cultured in RPMI 1640 + 10% FBS + 1% Penicillin-Streptomycin. All cells were maintained in a $CO_2$ incubator (Thermo Scientific, STERI-CYCLE i250) at 37 °C and 5% $CO_2$. Vero E6 cells (ATCC) were infected under biosafety level 3 conditions with SARS-CoV-2 (nCoV/Washington/1/2020, kindly provided by the National Biocontainment Laboratory, Galveston, TX).

### Supplemental Figures

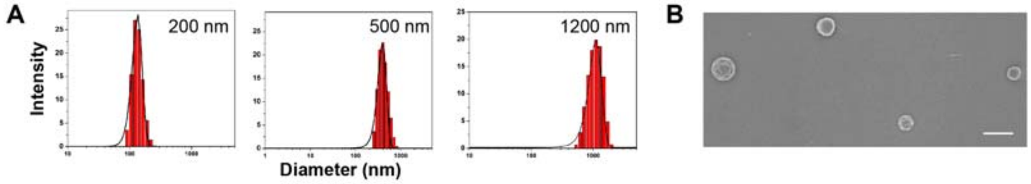


**Figure S1. Schematic design, synthesis, and characterization of Nanotraps for SARS-CoV-2**. (**A**) Size distribution of the Nanotraps with varying sizes measured by dynamic light scattering. (**B**) SEM image of dispersion of Nanotraps (500 nm) without virus. Scale bar represents 1 μm.

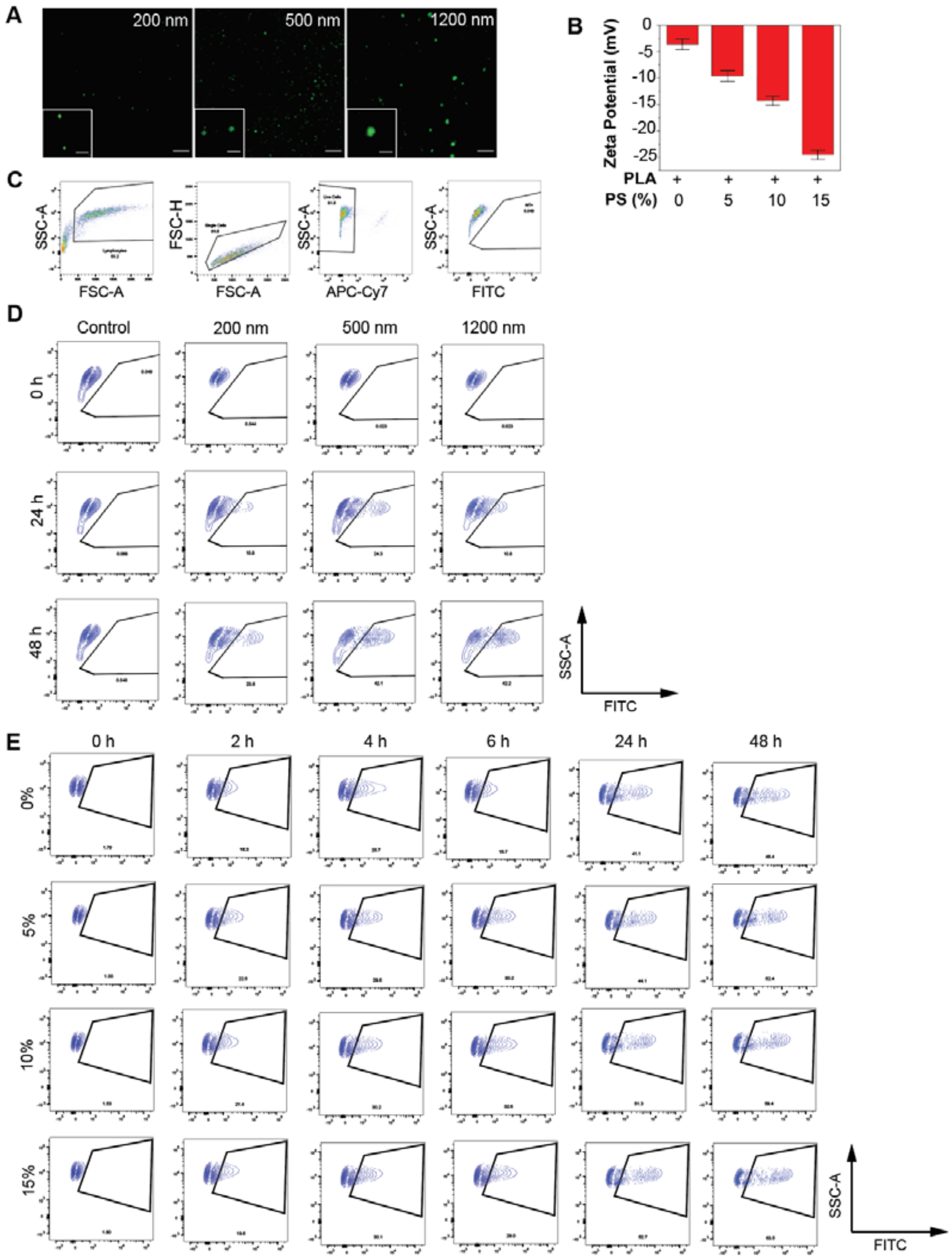


**Figure S2. Phagocytosis of Nanotraps by macrophages.** (**A**) Epifluorescent images of varying sizes of Nanotraps (DiO, green). (**B**) Measured zeta potentials of Nanotraps with varying PS ratios. (**C**) Gating strategy of the flow cytometry shown in Fig. 2aA-D. APC-Cy7 represents live/dead, and FITC represents DiO. (**D**) Contour plots of the flow cytometry shown in Fig. 2a-b. (**E**) Contour plots of the flow cytometry shown in Fig. 2c-d.

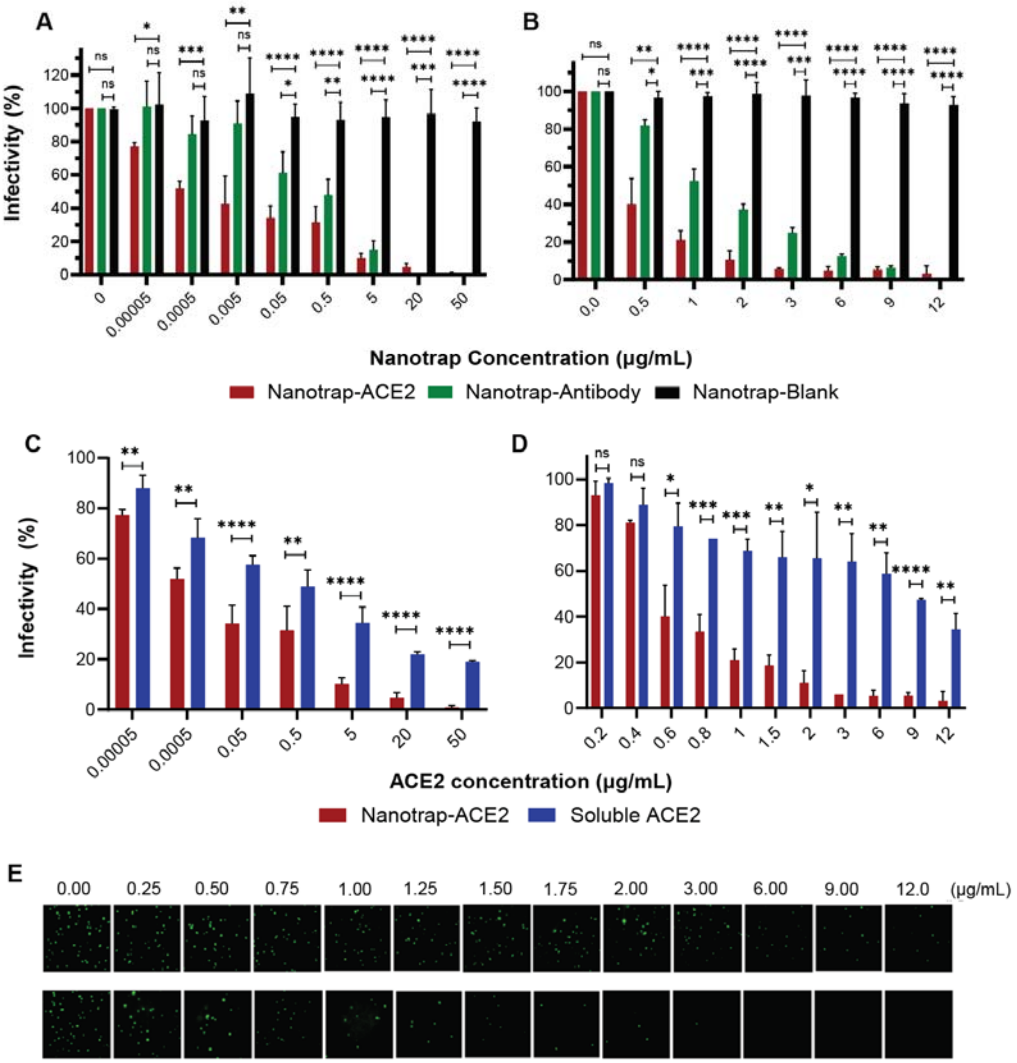


**Figure S3. Inhibition of SARS-CoV-2 pseudovirus in HEK293T-ACE2 cells**. (**A-B**) HEK293T-ACE2 cells were treated with SARS-CoV-2 spike pseudotyped lentivirus (A) or VSV (B) and Nanotrap-ACE2, Nanotrap-Antibody, or Nanotrap-Blank for 72 and 24 hours, respectively. Data are presented as mean ± SD; unpaired t tests were conducted on three independent replicates. (**C-D**) HEK293T-ACE2 cells were treated with SARS-CoV-2 spike pseudotyped lentivirus (C) or VSV (D) and Nanotrap-ACE2 or soluble ACE2 for 72 and 24 hours, respectively. Data are presented as mean ± SD; unpaired t tests were conducted on three independent replicates. (**E**) Representative fluorescent images of VSV-GFP pseudovirus infected HEK293T-ACE2 cells after treatment with ACE2 protein (top) and Nanotraps-ACE2 (bottom). *$p < 0.05$, **$p < 0.005$, ***$p<0.0005$, ****$p<0.00005$. N = 3 per group.

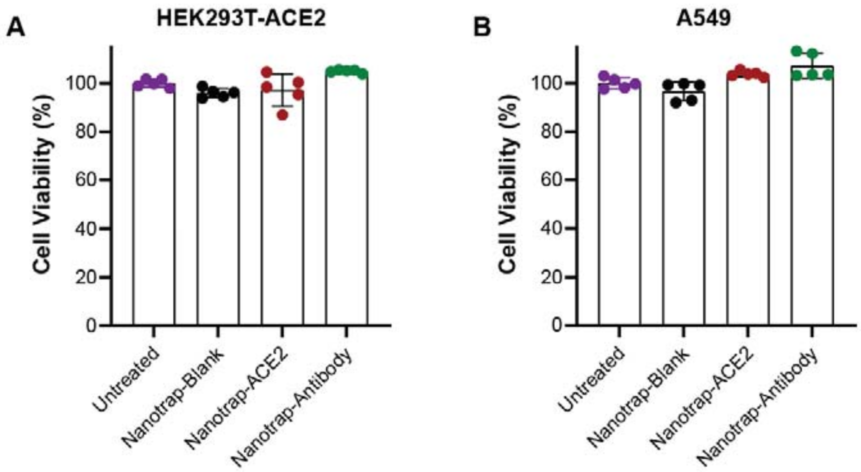


**Figure S4. *In vitro* biosafety profile of Nanotrap treatment.** (**A-B**) Viability of HEK293T-ACE2 (A) and A549 (B) cells with Nanotrap treatments for 72 hours. Data are presented as mean ± SD; n = 5 per group.

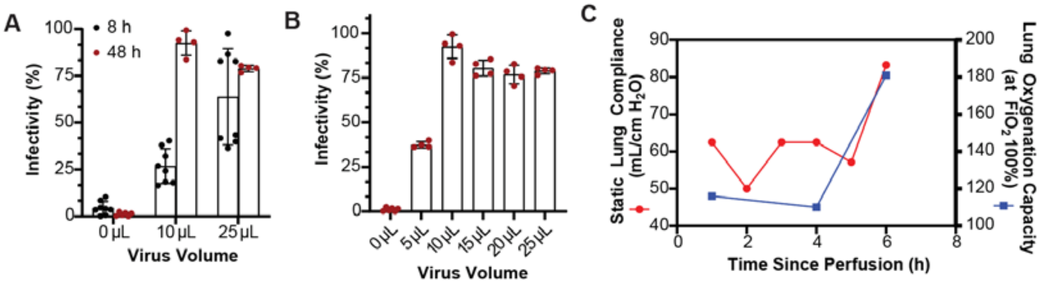


**Figure S5.** ***Ex vivo*** **human lung perfusion system for evaluating the neutralizing ability of the Nanotraps.** (**A**) SARS-CoV-2 pseudotyped lentivirus infectivity of primary human cells *in vitro* 8 or 48 hours post infection. (**B**) SARS-CoV-2 pseudotyped lentivirus infectivity of primary human cells *in vitro* 48 hours post infection. (**C**) Quantification of lung compliance and oxygenation capacity while on EVLP.

**Table S1. Formulations of lipid shell with different phosphatidylserine densities (molar ratios)**

| Formulation | DOPS | DSPC | Cholesterol | DSPE-mPEG$_{2000}$ | DSPE-PEG$_{2000}$-biotin |
|---|---|---|---|---|---|
| 0% PS | 0% | 65% | 30% | 4.5% | 0.5% |
| 5% PS | 5% | 60% | 30% | 4.5% | 0.5% |
| 10% PS | 10% | 55% | 30% | 4.5% | 0.5% |
| 15% PS | 15% | 50% | 30% | 4.5% | 0.5% |

### Supplemental Videos

**Supplemental Video 1: Lattice light-sheet microscopy imaging of macrophage phagocytosis.** 3-dimensional image reconstruction of macrophages (Wheat Germ Agglutinin (WGA)-CF488, green) phagocytosing Nanotraps (DiD, magenta). Video shows macrophages treated with 10% phosphatidylserine Nanotraps for 24 hours and untreated macrophages.

**Supplemental Video 2: Human *ex vivo* lung perfusion (EVLP) system.** Video shows lungs from **Figure 5** attached to the EVLP ventilator system.
